# Powassan Virus LB Neurovirulence and Lethality is Determined by Envelope Protein Domain III Residues

**DOI:** 10.64898/2026.03.26.714546

**Authors:** Marissa R. Lindner, Elena E. Gorbunova, Marcos Romario Matos de Souza, Grace E. Himmler, Hwan Keun Kim, Erich R. Mackow

**Author notes:** Corresponding Author Erich R. Mackow, Ph.D., Department of Microbiology and Immunology, Renaissance School of Medicine, Life Sciences Rm 126, Stony Brook University, Stony Brook, NY 11794-5222.

## Abstract

Powassan Viruses (POWV) are tick-borne Flaviviruses that cause lethal encephalitis and long-term neurologic sequelae. POWVs are divided into two lineages that share 86% genetic identity. POWV-LB (Lin I) was isolated from a fatal case of encephalitis, while POWV-LI9 (Lin II) is a tick-derived isolate that causes age-dependent lethality in mice. Domain III of Flavivirus envelope proteins (EDIII) directs cell attachment, and determines murine neuropathology and lethality. Here we developed POWV LB reverse genetics, analyzed the CNS of LB and recLB mutant infected mice, and assessed the role of EDIII residues in LB pathogenesis. LB infection of 50-week-old B6 mice focally activated microglia in the midbrain and cerebellum discrete from, and independent of, the location of envelope protein in the cerebral cortex, hippocampus and thalamus. Reverse genetic analysis of LB EDIII residues revealed that mutating D308N only partially reduced LB neuropathology and lethality. However, mutating D308N along with an adjacent EDIII residue, A310T, completely abolished LB-D308N/A310T neurovirulence, neuroinvasion and lethality, yet elicited neutralizing antibody responses that protected mice from a lethal WT LB challenge. Signifying changes in neurotropism, LB-D308N/A310T failed to activate microglia or yield detectable POWV RNA and antigen in the CNS. Overall, recombinant LB-D308N/A310T mutants fail to induce neurologic symptoms or neuropathology while eliciting a protective immune response. These findings reveal that EDIII residues determine the neurovirulence and lethality of lineage I POWVs and that genetically modifying EDIII provides a mechanism for POWV attenuation and vaccine development.

**Importance:** Powassan virus causes lethal encephalitis and long-term neurological sequalae in survivors; however, there are currently no licensed POWV vaccines or therapeutics. POWVs are comprised of two genetic lineages (LB-Lin I; LI9 Lin II) that are derived from discrete ticks and small mammal hosts. Flavivirus envelope proteins determine cell attachment, viral tropism, and neurovirulent spread and lethality in mice. Here we created a POWV LB reverse genetics system and assessed mutations that determine LB neurovirulence in 50 wk old mice. Studies reveal novel LB neuropathology with divergent patterns of gliosis and POWV antigen expression within the CNS. Mice infected with an LB-D308N/A310T envelope mutant elicited responses that protect mice from a lethal LB challenge and failed to enter the CNS, cause neurologic symptoms, CNS pathology or lethality. LB reverse genetics permits defining determinants of LB neurovirulence that are required for rational vaccine development.

## Introduction

Flaviviruses (FVs) are enveloped, positive-strand RNA viruses that are transmitted by tick and mosquito arthropod vectors worldwide (1, 2). Two tick-borne FVs, tick-borne encephalitis virus (TBEV) in Eurasia, and Powassan virus (POWV) in North America, cause a 10-15% fatal encephalitis that results in long term neurologic deficits in 50% of survivors (2, 3). POWV strain LB was isolated in 1958 from a 5-year-old with fatal encephalitis, and since then discrete POWVs have been isolated from expanding tick populations (4–7). Increased awareness and screening reveal more POWV seroprevalence than clinical cases, suggesting that many POWV infections are undiagnosed or asymptomatic (5, 6, 8). POWV is present in tick saliva injected into bite sites and this accounts for rapid POWV transmission, requiring as little as a 15-minute tick attachment (9, 10). POWV causes severe encephalitis, with cases rising with patient age and increasing severity and lethality in patients >49 years of age (2, 5, 7, 11–13). There are currently no licensed POWV vaccines or therapeutics.

Flaviviruses contain an ∼11 kb genome that encodes a polyprotein composed of 3 structural proteins (capsid, prM and envelope) and 7 nonstructural proteins (NS1/2A/2b/3/4A/4B/5) that are cleaved by cellular and viral proteases (2, 14). The viral surface is composed of 90 envelope protein dimers that form an icosahedral structure with 5-fold, 3-fold and 2-fold axes of symmetry that surround an internal lipid bilayer (15–17). The envelope protein is comprised of 3 domains (ED I-III) that are targets of neutralizing antibodies, direct viral attachment and fusion with cellular membranes and are associated with viral dissemination and virulence (18–20).

POWVs are divided into two genetically distinct lineages that are ∼86% identical at the nucleic acid level (21–23). Prototypic Lineage I strain LB, and Lineage II strains, Spooner and LI9, are sustained within discrete zoonotic cycles (23, 24). Lineage I POWVs are spread by *Ixodes marxi* and *Ixodes cookei*, with reservoirs including ground hogs, squirrels and skunks, while Lineage II POWVs are primarily spread by *Ixodes scapularis,* with its main reservoir being the white footed mouse (2, 24). Despite vector, host and genetic differences, Lineage I & II POWVs are highly conserved, sharing 96% identical envelope proteins, that form a single POWV serotype (21).

POWV LB was isolated by intracranial inoculation and passage in mouse brains prior to tissue culture adaptation (4). In contrast, POWV LI9 was isolated from deer ticks by the direct inoculation of VeroE6 cells (25). LB lytically infects cells, forming plaques in cell culture, while LI9 infection of VeroE6 cells is non-lytic, spreading focally cell-to-cell (25). In 50-week-old B6 mice LI9 causes microgliosis and spongiform lesions with ∼80% mortality 10-15 dpi (26). Surviving LI9 infected mice produce high level neutralizing antibody responses along with persistent CNS microgliosis (26, 27). LB causes lethal neurovirulent disease in mice resulting in meningitis, inflammatory infiltrates, perivascular cuffing and high CNS viral loads (28, 29). Whether initial intracranial isolation and passage of LB in mice contributes to lytic replication *in vitro* and novel neuropathology *in vivo* remains to be investigated.

Persistent passage of LI9 in VeroE6 cells produced an attenuated POWV strain, LI9P, with 8 amino acid differences from LI9 (27). Foot-pad inoculation of LI9P into 50-week-old B6 mice resulted in 100% survival without neurological sequelae, neuropathology or viral spread to the CNS (27). Flavivirus cell attachment and virulence is linked to envelope protein domain III (EDIII), and attenuated LI9P contains a single EDIII D308N mutation (27, 30–32). Although additional LI9P mutations impact virulence, mutating just the D308N residue in LI9 (LI9-D308N) mirrored LI9P attenuation, and failed to cause neurologic symptoms, neuropathology or lethality in aged B6 mice (27).

LB differs genetically and phenotypically from LI9 *in vitro* and *in vivo*. The source of functional differences remains unknown but likely stems from the 3-10% dissimilarity of LB and LI9 structural and nonstructural proteins. Here we examined the role of envelope protein EDIII residues in POWV LB neurovirulence, neurotropism and lethality in mice. To accomplish this, we generated a circularized polymerase extension reaction (CPER) reverse genetics system for POWV-LB that robustly produces infectious recombinant LB and LB-EDIII mutants. Inoculation of recLB into 50-week-old B6 mice resulted in weight loss, neurologic sequelae and 79% lethality. In contrast to LI9, mutating D308N in POWV LB only delayed neurologic sequelae and reduced lethality (33%) but failed to completely attenuate LB. Analysis of LB and LI9 envelope proteins revealed an alanine to threonine substitution in EDIII adjacent to D308, that was targeted for mutagenesis. Mice inoculated with the double EDIII mutant, LB-D308N/A310T, abolished LB directed neurological symptoms, microglial activation, CNS histopathology and lethality in 50-week-old B6 mice. In contrast, the CNS of mice infected with WT LB revealed predominant envelope protein immunostaining in the cerebral cortex independent of focal microglial/macrophage activation in the midbrain. Overall, our studies establish a lineage I POWV LB reverse genetics system and demonstrate a fundamental role of envelope protein EDIII residues in determining POWV LB neurovirulence.

## Results

### POWV LB Reverse Genetics

The age-dependent severity of human POWV encephalitis is mirrored by the infection of B6 mice with LI9, a Lineage II POWV isolate (26). Using reverse genetics, D308N mutations introduced into the LI9 envelope protein blocked neurologic symptoms, CNS pathology and lethality in B6 mice (27). Prototypic Lineage I POWV, LB, also causes lethal encephalitis in humans and mice, however determinants of LB neurovirulence have yet to be defined, and reverse genetics systems for analyzing LB mutants remain to be developed (4, 28, 29, 33, 34). Lineage I and Lineage II POWVs share 86% nucleotide identity (Fig. 1A), yet envelope proteins of LB and LI9 are 96% identical (97% similar) and comprise a single serotype (21). Envelope protein conservation suggests the potential for common LB and LI9 envelope residues to direct CNS neurovirulence and lethality. In order to evaluate determinants of LB pathogenesis, we devised a reverse genetics system compatible with LB RNA sequences. Using LB-specific oligos (Table 1), five LB fragments (∼2-2.2 kb each) with 26-nucleotide overlapping sequences were cloned into pMiniT plasmids that cover the ∼11 kb LB genome (Fig. 1B). To circularize LB fragments and direct RNA transcription, a sixth overlapping clone (530 bp) was generated, containing a CMVd2 promoter, SV40 polyadenylation site and Hepatitis Delta Virus ribozyme (HDVr) (Fig. 1C) (27, 35). Fragments connected by a circular polymerase extension reaction (CPER) were transfected into BHK-21 cells and 3 dpt cell supernatants were used to infect VeroE6 cells (Fig. 1D). RecLB stocks were passaged in VeroE6 cells (P1-P4), sequence verified and observed to focally infect cells, with a lytic phenotype analogous to WT POWV LB (Fig. 1E).

**Figure 1.**
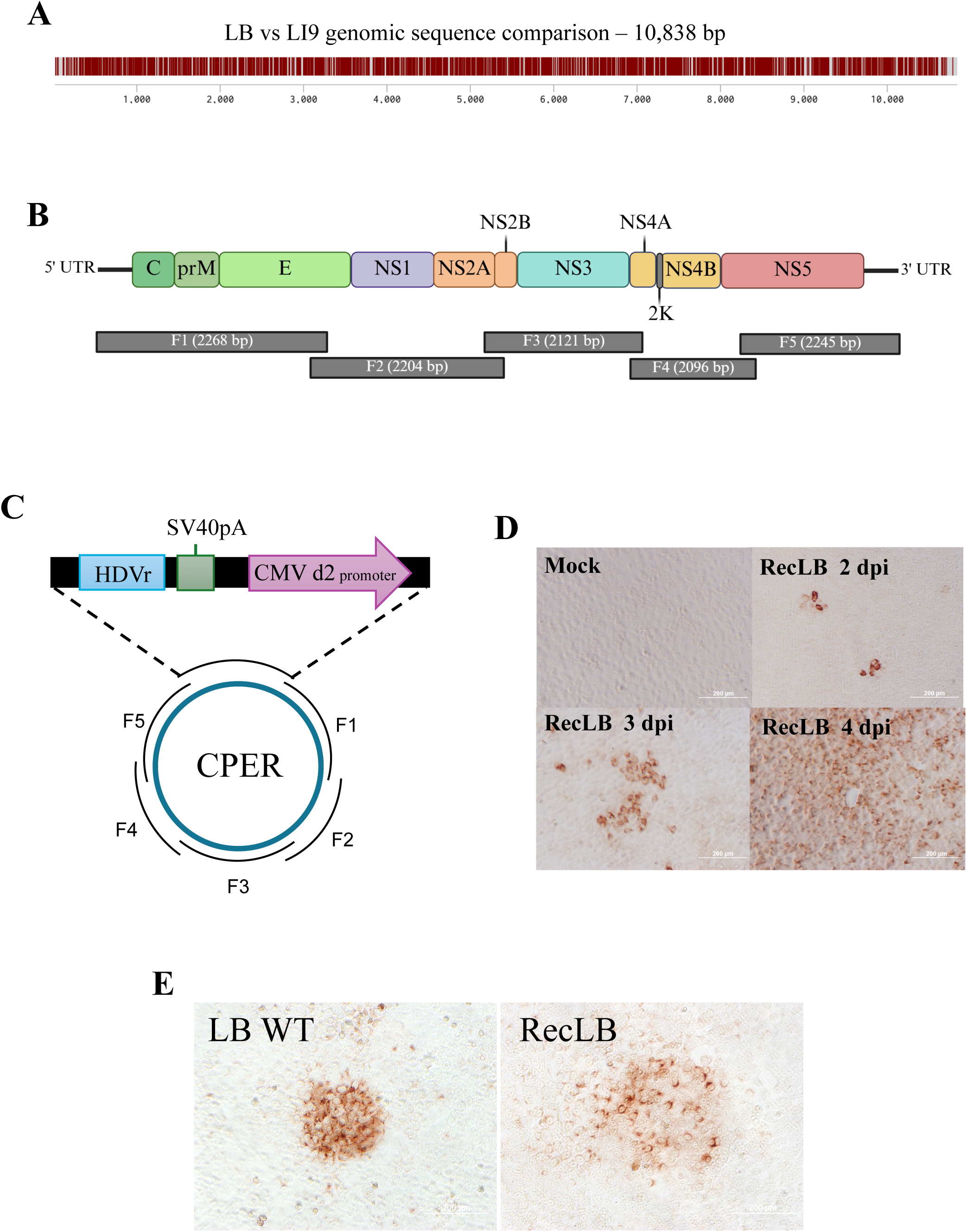
Lineage I POWV LB CPER Reverse Genetics System. (A) Genetic homology of POWV strains LB and LI9 with differences indicated by white bars. (B) LB genomes (∼11 kb) were cloned into 5 overlapping fragments and purified Phusion amplified fragments were used develop reverse genetics. (C) POWV-LB fragments and UTR linkers, containing an HDVr, SV40pA, and CMVd2 promoter, were CPER circularized prior to transfection. (D) Immunoperoxidase staining of recLB infected VeroE6 cells 2-4 dpi. (E) Immunoperoxidase staining of WT LB or recLB infected VeroE6 cell foci (MOI, 0.1) 4 dpi.

**TABLE 1.**
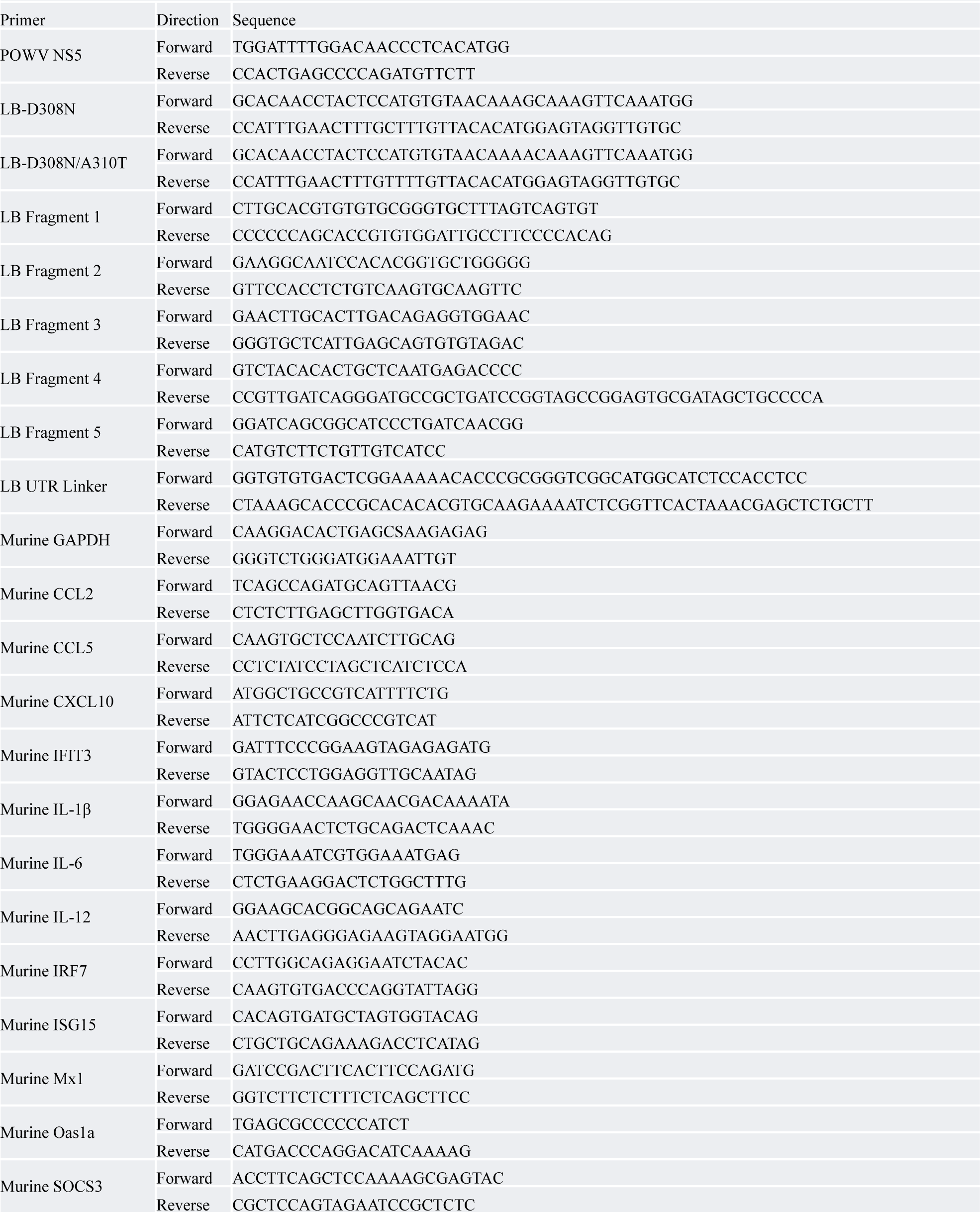
Primers.

### Generation of POWV LB Envelope Protein Mutants

We previously defined LI9 EDIII mutations as critical determinants of neurovirulence and lethality in B6 mice (27). LI9 and LB envelope proteins are 96% identical with EDIIIs that differ by just a single T310A residue between cysteines 307 and 399 (Fig 2A). Pymol visualized LB and LI9 EDIII structures (36) highlight the location of D308 and A310T residue differences in the context of a previously defined D308 to K311 salt bridge (37)(Fig 2B). In order to define roles for EDIII residues in LB neurovirulence and lethality, we generated LB mutants with single (LB-D308N) or double (LB-D308N/A310T) EDIII residue changes. EDIII mutants (Fig 2A) were generated by site-directed mutagenesis of Fragment 1 clones in pMiniT plasmids (Table 1). Mutated EDIII Fragment 1 was amplified, CPER assembled with F2-F5 and linker fragments and transfected into BHK-21 cells. RecLB-EDIII mutant stocks were sequence verified and shown to spread focally and replicate with similar kinetics and titers to WT recLB 2-4 dpi (Fig 2C,D).

**Figure 2.**
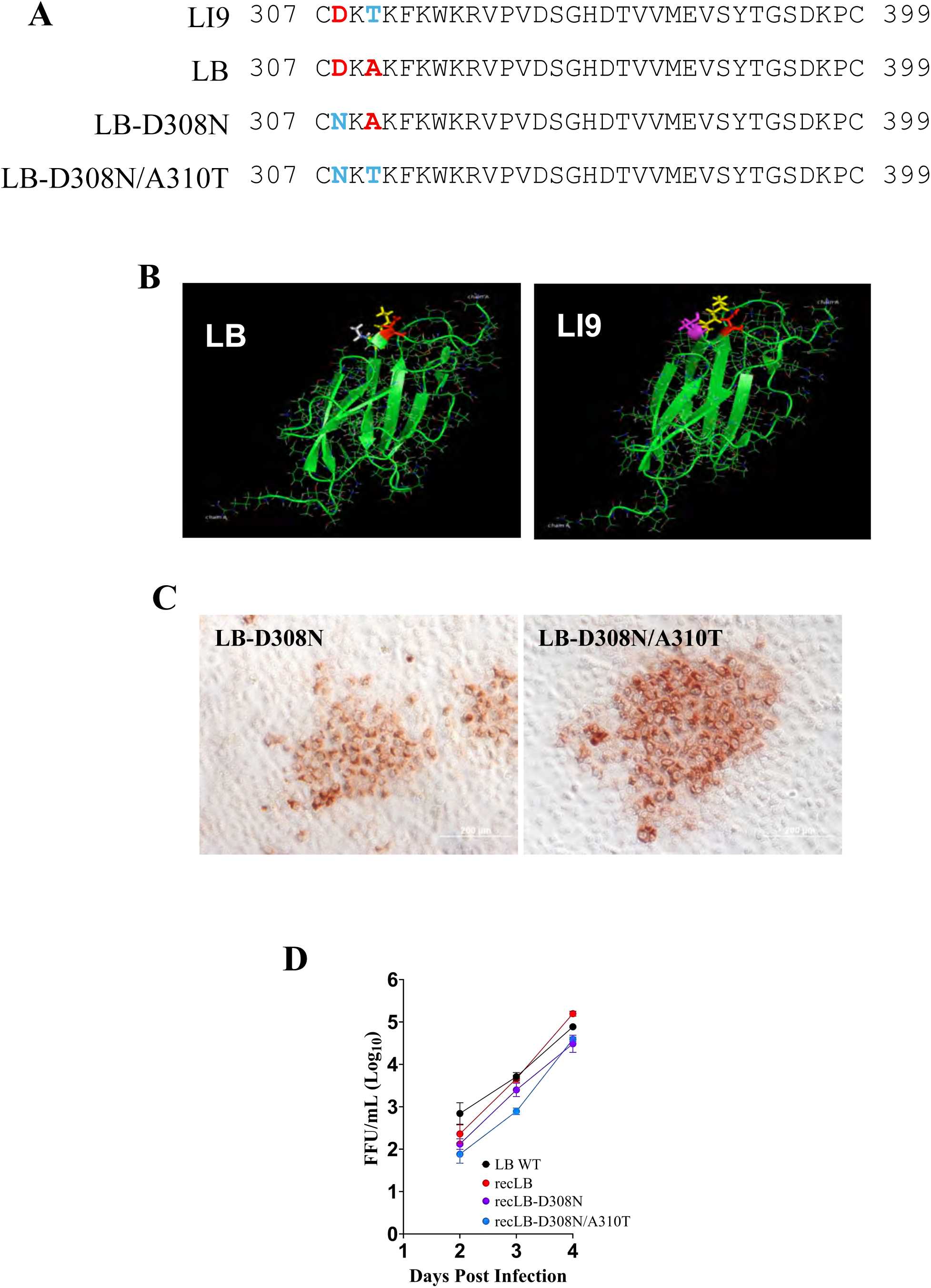
Generation of LB EDIII mutant viruses. (A) Alignment of LB and LI9 EDIII residues between C307 and C399 and CPER generated LB mutants. (B) LB and LI9 POWV Envelope domain III structures (Pymol) comparing LB (left) and LI9 (right) with highlighted LB A310 (white), LI9 T310 (pink) and D308 (red) - K311 (yellow) putative salt bridge residues. (C) VeroE6 cells infected with CPER derived LB-D308N and LB-D308N/A310T mutants (MOI, 0.1) were immunoperoxidase stained with anti-POWV HMAF 4 dpi. (D) VeroE6 cells were LB, LB-D308N, or LB-D308N/A310T infected (MOI, 0.1) and viral titers in cell supernatants were assayed in duplicate 2-4 dpi. Data is presented as the mean ± SD and was analyzed via two-way ANOVA.

Similar to mutating LI9-D308N, the LB-D308N/A310T double mutant results in the generation of a potential N-linked glycosylation site (NxT) in the envelope protein. However, in both envelope proteins NxT sites are adjacent to cysteine residues with intervening lysine residues (CNKT) that normally restrict N-linked glycosylation. We previously found no additional glycosylation of LI9-D308N envelope protein, and analogously evaluated whether LB-D308N/A310T contains additional glycosylation. The electrophoretic migration of the envelope protein expressed from WT LB and LB-D308N/A310T infected cells was compared by Western blot (Fig. S1). We found no decrease in the electrophoretic mobility of the LB-D308N/A310T mutant envelope protein relative to the WT envelope protein, indicating that the CNKT motif in the LB-D308N/A310T mutant does not appear to be a viable glycosylation site (Fig. S1).

### EDIII Mutations Determine the Lethality of POWV LB in B6 Mice

To investigate whether LB neurovirulence is determined by EDIII residues we footpad inoculated 50-week-old C57BL/6 mice with 2000 FFU of LB, LB-D308N, or LB-D308N/A310T viral mutants or PBS. LB infected mice began to lose weight 5 dpi (Fig. 3B) and became progressively lethargic and weak with ruffled fur prior to neurologic symptoms. LB infection of B6 mice was 79% lethal (N=19) with signs of neurological disease (weakness, twitching, and partial hind limb paralysis) and lethality occurring between 8 and 16 dpi (Fig. 3B,E). In contrast, LB-D308N inoculated mice (N=21) were partially attenuated with 7 mice experiencing rapid weight loss (Fig. 3C) that coincided with 33% lethality (Fig. 3E). In LB-D308N infected mice, lethality was delayed to between 12 and 25 dpi (Fig. 3E), with all 7 moribund mice showing neurological symptoms including weak grip and partial or total hindlimb paralysis. Despite this, in 14/21 mice (67%) LB-D308N infection failed to cause weight loss, signs of neurological disease or lethality (Fig. 3C, E). Sequencing viral RNA from the brains of moribund mice confirmed the presence of the LB-D308N mutant in the CNS and that neurovirulence was not the result of reversion to WT LB. In contrast, foot-pad inoculation of B6 mice with LB-D308N/A310T failed to cause weight loss (Fig. 3D) or neurological symptoms and resulted in 100% survival of B6 mice 30 dpi (N=21) (Fig. 3E). These findings demonstrate that both D308N and A310T EDIII residues are required for POWV LB attenuation, and that EDIII A310 residues are novel determinants of POWV LB neurovirulence and lethality.

**Figure 3.**
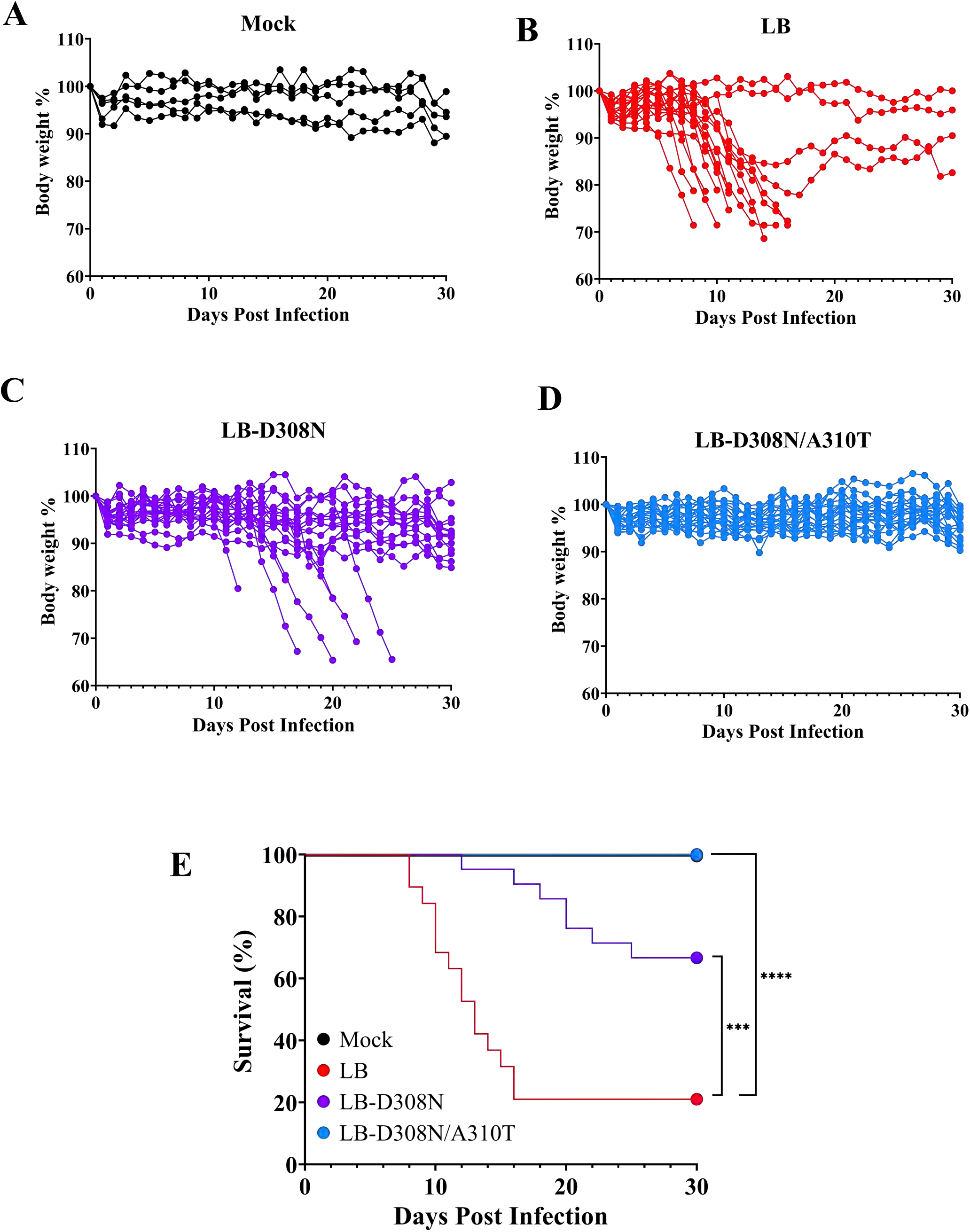
EDIII mutagenesis of LB attenuates lethal infections. Fifty-week-old C57BL6 mice were subcutaneously footpad inoculated with 2,000 FFU of LB (N=19), LB-D308N (N=21), LB-D308N/A310T (N=21), or mock (PBS)(N=5). Mice were weighed daily for 30 days following infection and assessed for clinical neurological signs of disease to determine humane lethal endpoints. (A-D) Individual mouse weights of Mock (A), LB (B), LB-D308N (C), and LB-D308N/A310T (D) infected mice are presented as a percent of their original bodyweight 0 dpi. (E) Kaplan-Meier curves of lethal POWV infection of LB, LB-D308N, LB-D308N/A310T, and mock were analyzed by log-rank test (\*\*\**P*<0.001; \*\*\*\**P* <0.0001). (A-E) Data are cumulative from two independent experiments.

### CNS Responses and Histopathology Following LB-D308N/A310T Infection

Mice infected with LI9, but not attenuated LI9P or LI9-D308N mutants, demonstrate dramatic increases in activated microglia (Iba1^+^), spongiform lesions and high viral loads in the CNS (26, 27). In contrast to LI9, background levels of viral RNA are observed in the CNS of LI9P and LI9-D308N infected mice suggesting that attenuation is coincident with restricted POWV neuroinvasion (27). To evaluate differences in LB and LB-D308N/A310T neuroinvasion, we assessed viral RNA levels in the CNS of 50-week-old mice by qRT-PCR (moribund, 10 dpi or 30 dpi). Consistent with neurovirulence and lethality, moribund LB infected mice had high levels of viral RNA in the CNS (∼10^5^ FFU equivalents/gm)(Fig. 4A). In contrast, CNS RNA levels were at background levels in LB-D308N/A310T infected mice (Fig. 4A) consistent with 100% survival and a lack of neurologic symptoms. In mice surviving LB or LB-D308N/A310T infection, analysis of CNS viral loads 30 dpi revealed no evidence of viral RNA over background levels, indicating that virus is cleared from the CNS after acute infection and absent in the CNS of survivors (Fig. 4A).

**Figure 4.**
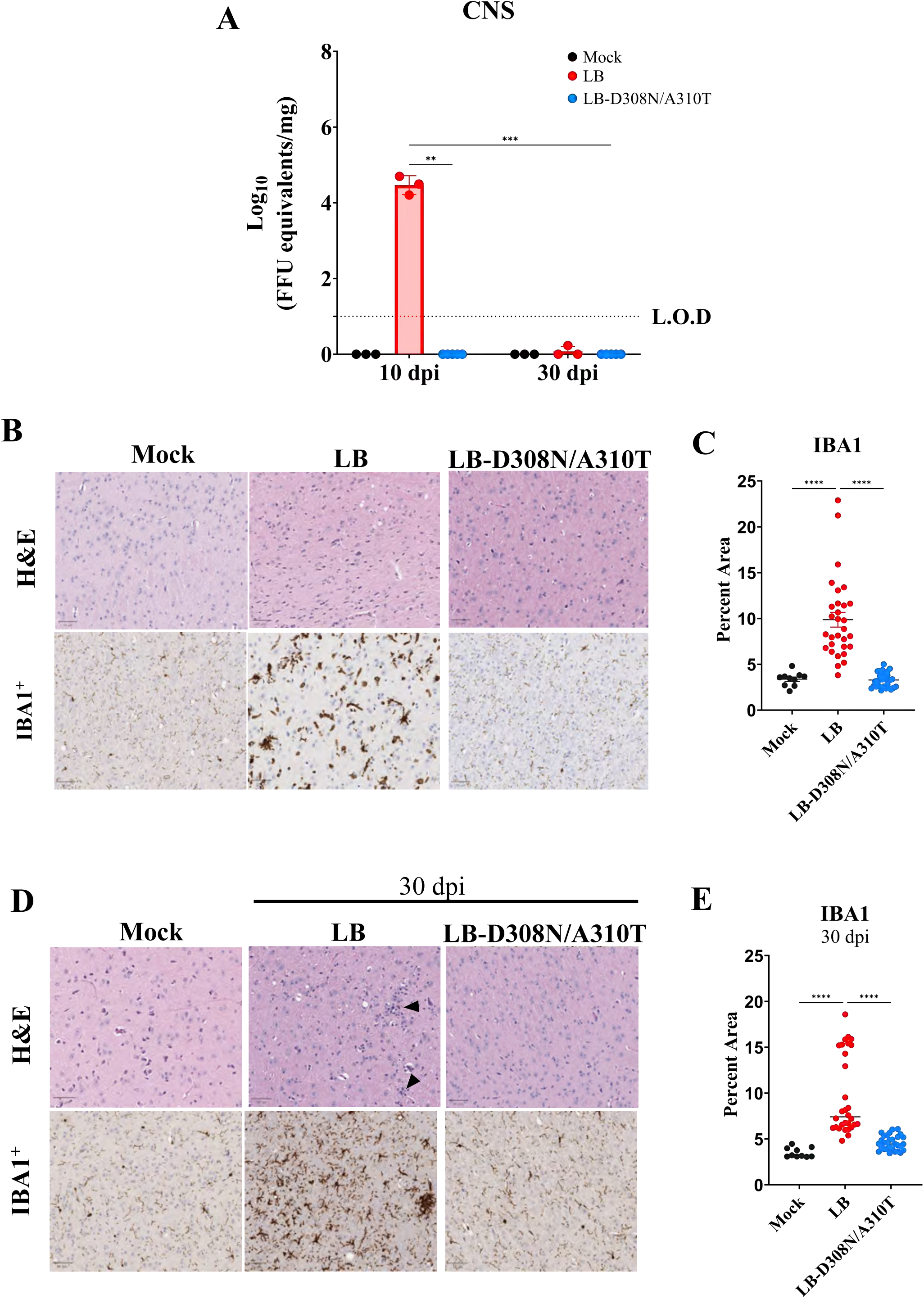
LB-D308N/A310T fails to activate CNS microglia or induce pathology. C57BL/6 mice (50-week-old) were subcutaneously inoculated with 2000 FFU of LB (N=3) LB-D308N/A310T (N=5), or mock infected (N=3). At 10 dpi (A, B, C) or 30 dpi (A, D, E), mice were euthanized and brains were either harvested for total RNA extraction (A) or fixed for immunohistochemical analysis (B,D). (A) Total RNA levels in the CNS of mice were assayed in duplicate by qRT-PCR and data expressed as the Log_10_ of POWV FFU equivalents per mg of tissue (mean ± SD) normalized to a POWV RNA standard curve. Data are representative of two independent experiments. Statistical analysis was determined by two-way ANOVA of log_10_ transformed values (\**P* < 0.05, \*\**P* < 0.001, \*\*\**P* < 0.0001). (B, D) Brains were sectioned, H&E stained and immunostained with anti-Iba1 for microglial activation. Representative images of the midbrains of mock, LB and LB-D308N/A310T infected mice are presented. Black arrowheads denote immune cell infiltrates in LB infected midbrains. ImageJ quantification of Iba1^+^ immunostaining reflects the percent Iba1^+^ pixel intensity from 10 midbrain regions of mice (N = 3) 10 dpi (C) or 30 dpi (E) vs mock (N=1). Data are presented as mean of individual quantitated regions and statistical significance was determined by one-way ANOVA (\*\*\*\**P*<0.0001).

Long term neurological sequalae is observed in ∼50% of patients that survive POWV infection (5, 11, 38, 39). To investigate CNS pathology, we analyzed the CNS of B6 mice infected with LB, or LB-D308N/A310T by hematoxylin and eosin (H&E) and anti-Iba1 immunostaining 10 or 30 dpi. Similar to mock infected CNS controls LB-D308N/A310T infected mice revealed no signs of aberrant histopathology or Iba1^+^ microglial/macrophage activation (Fig. 4B,C). In contrast, LB infected mice had evidence of spongiform lesions and dramatically increased Iba1^+^ staining (Fig. 4B,C). Analysis of murine survivors 30 dpi also revealed quantifiably higher numbers of Iba1^+^ staining in the CNS of LB, but not LB-D308N/A310T, infected mice (Fig. 4D,E). POWV envelope protein immunostaining of LB and LB-D308N/A310T infected CNS at 30 dpi revealed no signs of antigen within the CNS (Fig. S2). Taken together analysis of the CNS of LB survivors 30 dpi reveals residual Iba1^+^ inflammatory responses that persist despite clearance of virus from the CNS (Fig 4 A,D,S2). Overall, these data are consistent with failed LB-D308N/A310T neuroinvasion and suggest the virus independent persistence of CNS inflammation in LB survivors 30 dpi.

### Divergent Localization of POWV LB Antigen and Iba1^+^ Responses

Sagittal sections from the brains of B6 mice infected with LB, LB-D308N/A310T, or mock infected were immunostained for POWV envelope protein and Iba1^+^ positive microglial/macrophages. Consistent with a lack of RNA in the CNS of LB-D308N/A310T infected mice, envelope protein and Iba1 staining were identical in the CNS of mock or LB-D308N/A310T infected mice (Fig. 5). In contrast, LB infection resulted in prominent envelope protein staining in the cerebral cortex, thalamus, and hippocampal pyramidal neurons in the CNS (Fig. 5,6). Discrete from envelope protein localization, Iba1^+^ microglia/macrophage staining was minimal in the cerebral cortex and instead focally dispersed throughout the midbrain, cerebellum and pons (Fig. 5,6). Additional Iba1 staining was identified in the meninges, areas of perivascular cuffing and focal areas within the midbrain that are consistent with extensive inflammatory immune cell recruitment (Fig. 6). The discernable lack of coincident Iba1 and envelope staining in the cerebral cortex and thalamus is further observed in predominantly Iba1 positive areas of the medulla, midbrain, cerebellum and pons (Fig. 6). These findings reveal an inverse correlation between the location of POWV antigen and activated microglial/macrophage responses within the CNS, that are critical to understanding mechanisms of LB directed neuropathology.

**Figure 5.**
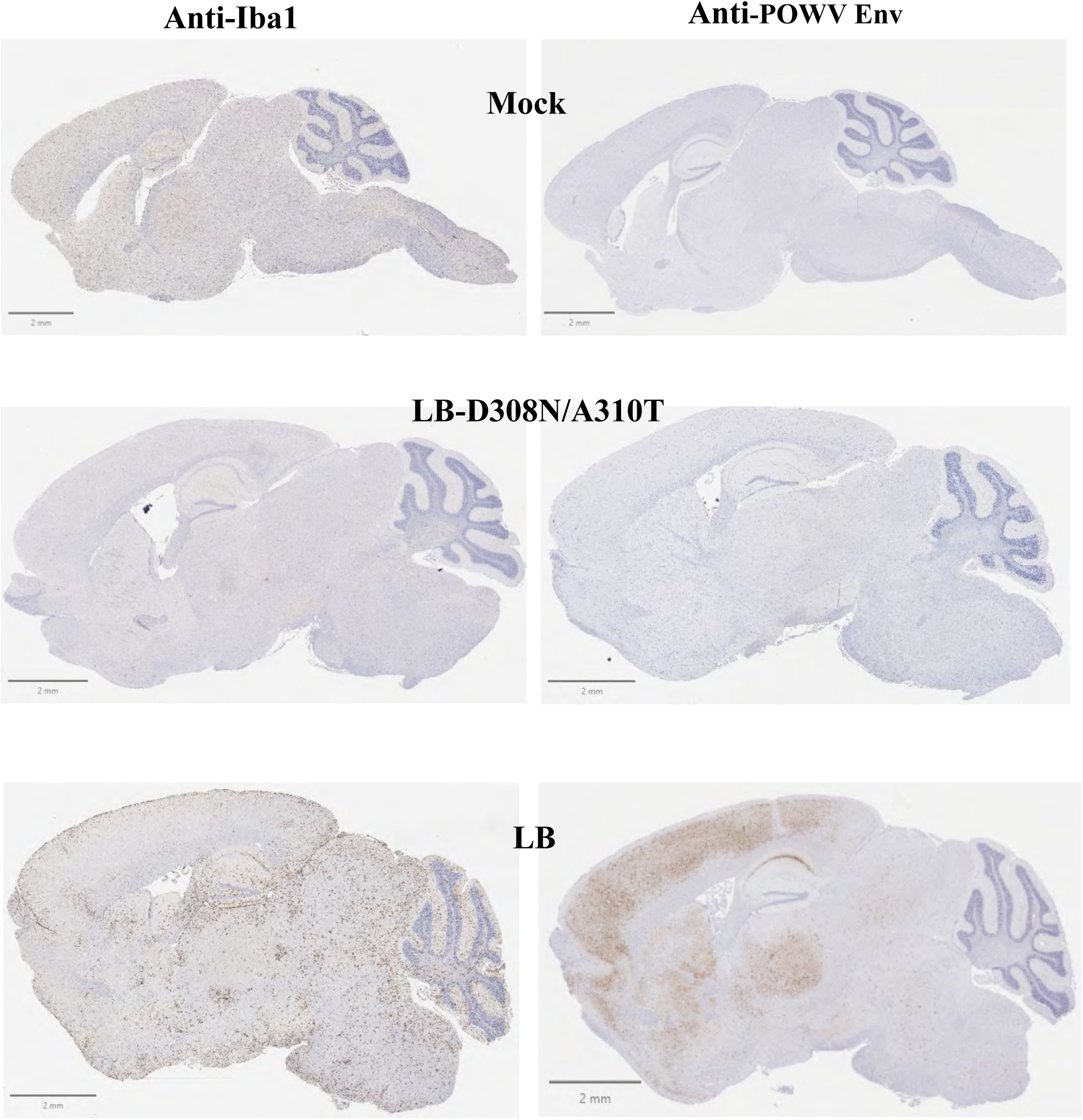
LB vs LB-D308N/A310T CNS infection. LB (N=3), LB-D308N/A310T (N=3) or mock (PBS) (N=1) infected 50-week-old C57BL/6 mice (2000 FFU) were euthanized 10 dpi (LB-D308N/A310T) or when moribund (LB) and brains were harvested, fixed and anti-Iba1 or anti-POWV envelope protein immunostained. Representative immunostaining of whole brain slices from LB and the LB-D308N/A310T mutant are presented.

**Figure 6.**
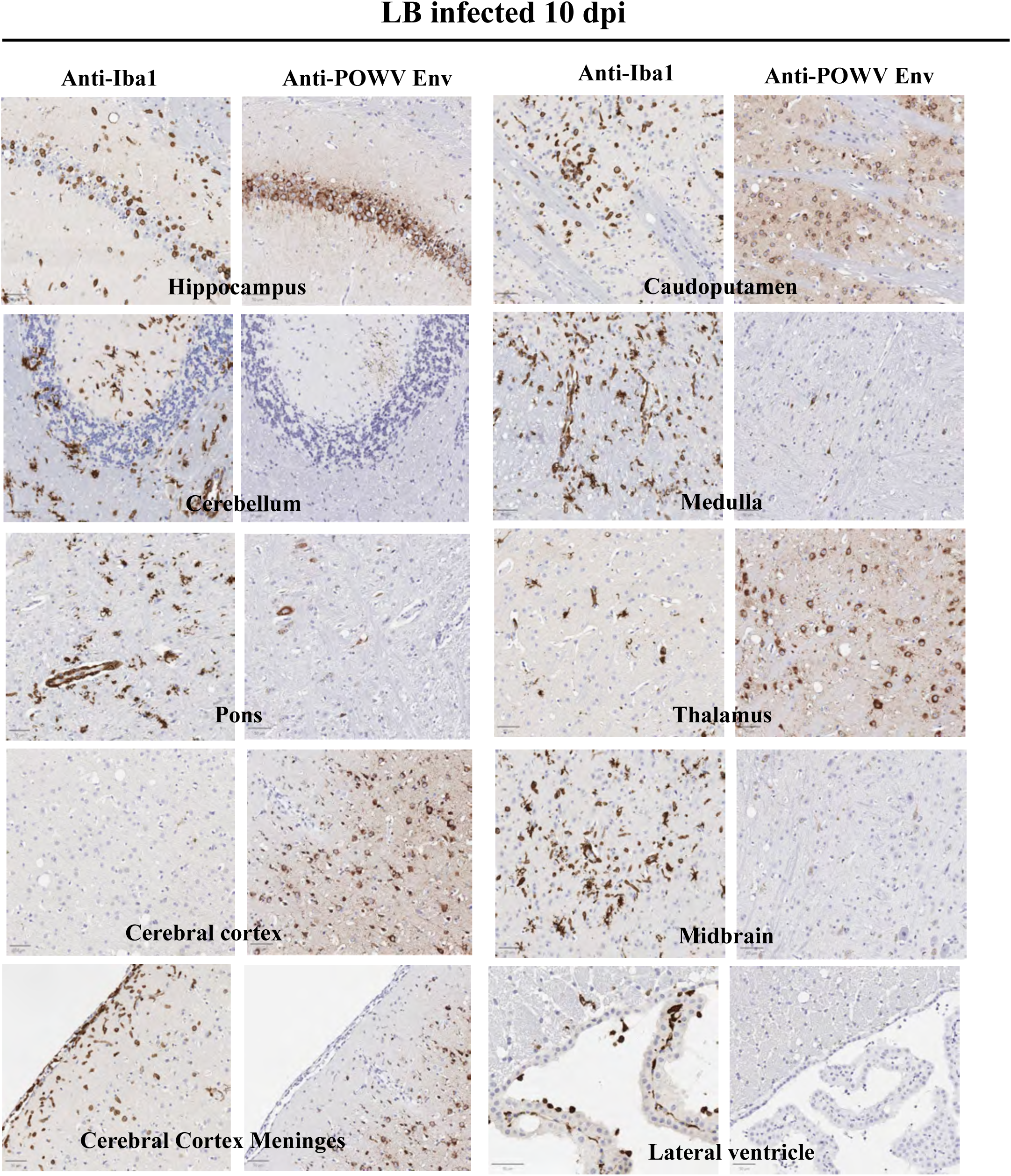
Distinct LB gliosis phenotype. Representative images from sequential sections of a 10 dpi LB infected mouse brain immunostained with anti-Iba1 (left) and anti-POWV envelope protein (right) are presented side-by-side from the hippocampus, cerebellum, pons, cerebral cortex, Cerebral cortex meninges, caudoputamen, medulla, thalamus, midbrain and lateral ventricle.

### LB, but not LB-D308N/A310T, Activates CNS Immune Responses

To compare LB and LB-D308N/A310T CNS responses, we assessed cytokine, chemokine and interferon stimulated gene (ISG) induction in the CNS by qRT-PCR. All assayed ISGs (IRF7, Mx1, IFIT3, SOCS3, ISG15, and OAS1a) were induced 10 to 400-fold following LB infection of mice, and at mock infected control levels in LB-D308N/A310T infected mice (Fig. 7A). Similarly, proinflammatory CXCL-10, CCL2, and CCL5 chemokines were induced ∼800 to 2000-fold in LB infected brains, while at mock infected control levels in the LB-D308N/A310T infected CNS (Fig. 7B). Proinflammatory cytokines IL-1β, IL-6, and IL-12 were also induced ∼30 to 1000-fold, respectively, in the CNS of LB infected mice but not in LB-D308N/A310T infected mice (Fig. 7B). Collectively, these results indicate that LB-D308N/A310T fails to induce proinflammatory and innate immune CNS responses and are consistent with failure of attenuated LB-D308N/A310T to infect the CNS or cause neurovirulent disease.

**Figure 7.**
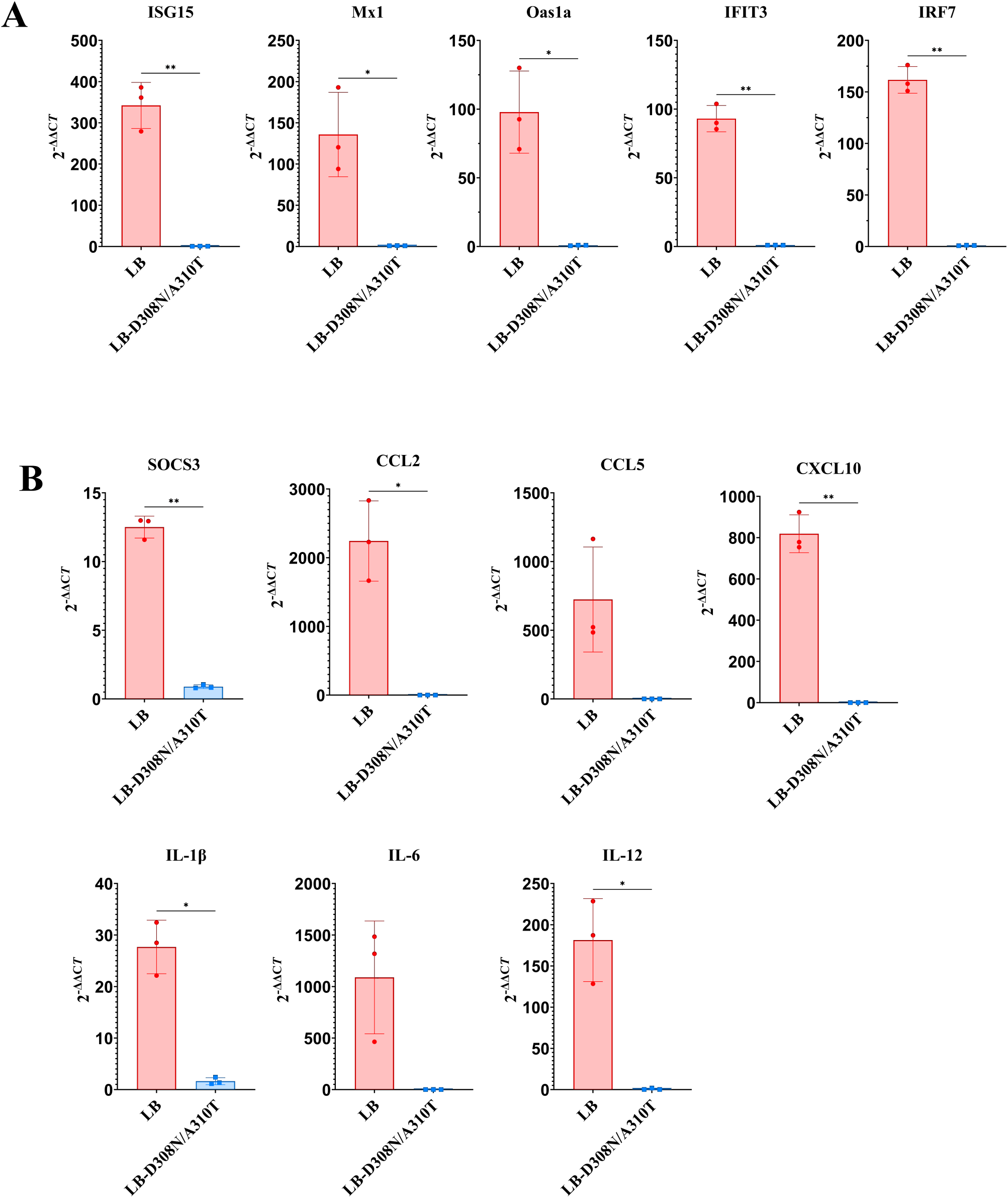
LB, but not LB-D308N/A310T, Induced CNS Responses. Total RNA was extracted from brains of the LB, LB-D308N and mock infected mice and the induction of (A) ISG, and (B) cytokine, and chemokine transcripts were analyzed by qRTPCR. Data is presented from one of two independent experiments as the 2^-ΔΔCT^, with each dot representing an individual mouse (N=3 per group). Statistical significance was determined by Welch’s T test (\**P*<0.05, \*\**P*<0.01).

### LB-D308N/A310T Inoculation Protects Mice from a Lethal LB Challenge

Here we determined if attenuated LB-D308N/A310T protects mice from a lethal LB challenge. Mock (N=5), or LB D308N/A310T (N=9) infected 50-week-old B6 mice were retro-orbitally bled 28 dpi and challenged on day 30 with a lethal dose of WT LB (2000 FFU). LB-D308N/A310T infected mice produced high titer LB neutralizing antibodies (1:160-1:640). Mock-infected mice challenged with LB mice lost weight, showed neurological symptoms, with 80% succumbing to LB infection (Fig. 8B,D). Mice previously infected with LB-D308N/A310T failed to lose weight or display neurological symptoms and 100% survived the lethal LB challenge (Fig. 8C,D). These findings reveal that LB-D308N/A310T infection elicits a protective immune response and indicate that mutating LB-EDIII residues is a viable mechanism of POWV LB attenuation and vaccine development.

**Figure 8.**
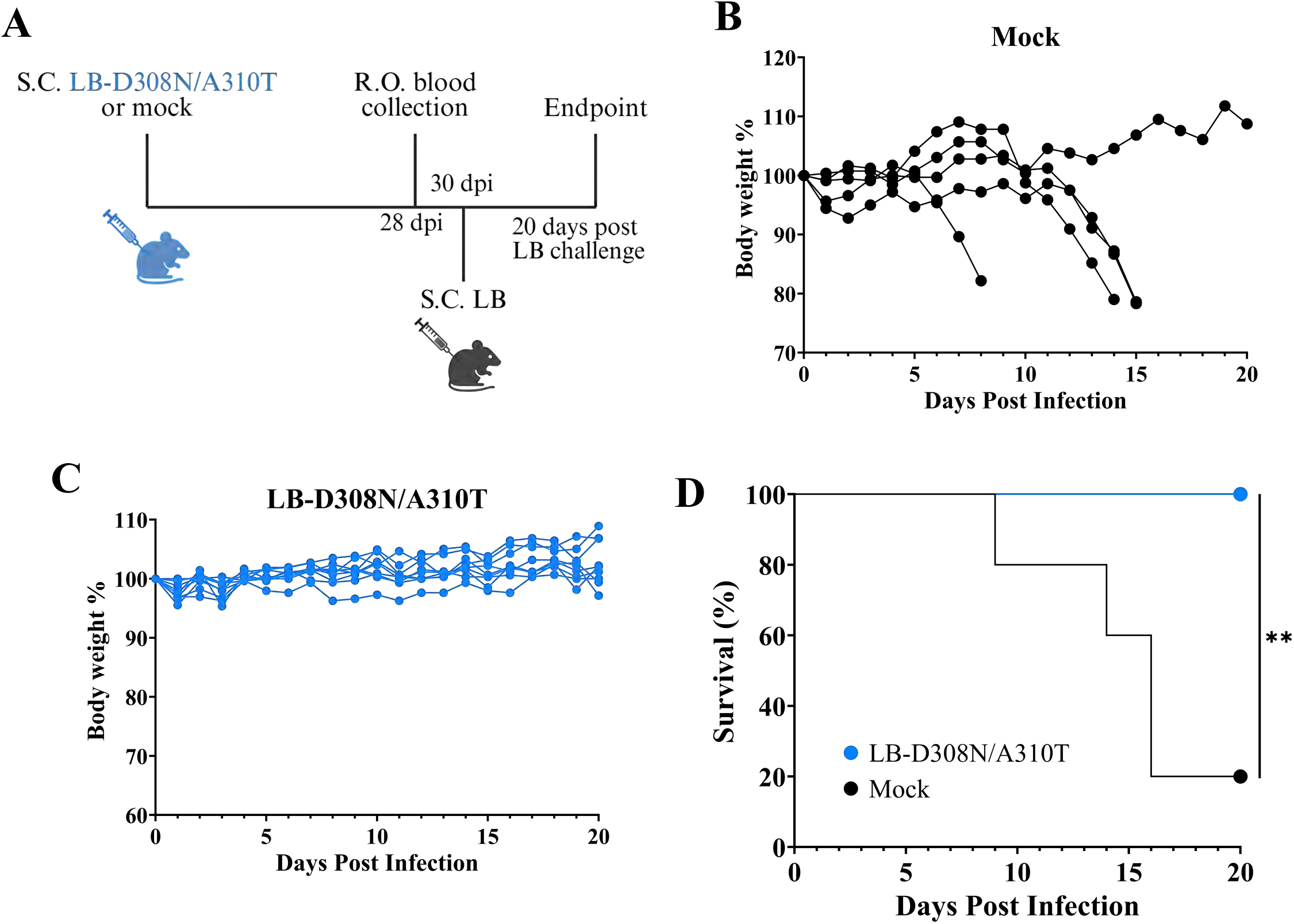
LB-D308N/A310T Inoculate Mice are Protected from a Lethal LB challenge. (A) Experimental timeline in days. Mice were infected subcutaneously with 2000 FFU of LB-D308N/A310T (N=9) or mock infected (PBS) (N=5). All mice survived initial infections without weight loss or neurological symptoms. At 28 dpi mice were retro-orbitally bled and at 30 dpi challenged with 2000 FFU of LB. Mice were monitored daily for changes in body weight and clinical neurological symptoms. (B, C) Individual mouse body weights were monitored 20 dpi after challenge and presented as percent of original body weights prior to the lethal LB challenge on 30 dpi. (D) Kaplan-Meier curves of lethal LB challenged mice (LB-D308N/A310T and naïve survivors) were analyzed by log-rank test (\*\**P<0.01*).

## Discussion

POWVs cause lethal encephalitis and long-term neurologic deficits with disease severity and lethality increased in individuals >49 years of age (2, 5, 7, 40). Despite age-dependent incidence, POWV infections are diagnosed across age groups and prototypic POWV LB was isolated from the brain of a 5 year old with fatal encephalitis (4). LB was passaged in mouse brains and causes lethal disease in mice independent of age, while tick-derived LI9 causes age-dependent lethality in mice, mirroring human POWV severity (4, 11, 26, 29). LB and LI9 direct lethal encephalitis in immunocompetent B6 mice without IFN inhibition required for some Flaviviruses (26, 28, 41, 42), although mechanisms of POWV neuroinvasion and neurovirulence remain enigmatic.

Flavivirus envelope proteins are targets of neutralizing antibodies, and associated with cell attachment, tropism and virulence (18–20, 30–32, 43–46). EDIII forms an immunoglobulin-like structure at the 5-fold axis of symmetry on the surface of FVs that is associated with viral binding to cell surface receptors (15, 37, 47, 48). Although POWV receptors have yet to be defined, EDIII is presumed to direct POWV cell attachment, dissemination and neurotropism, and prior findings link EDIII residues of POWV LI9 to lethal neurovirulence in a murine model (18, 27, 46, 47, 49). Passage of WT LI9 POWV in VeroE6 cells yielded LI9P, an attenuated strain containing an EDIII D308N mutation, that lacks neurovirulence and lethality in mice (27). Inserting just D308N into WT LI9 abolished the lethality and neurovirulence of LI9-D308N, while eliciting responses that protect mice from a lethal LI9 challenge (27). In fact, attenuated LI9P or LI9-D308N POWVs fail to enter the CNS or activate CNS responses, suggesting that EDIII mutations prevent LI9 neuroinvasion (27).

POWV LB differs from LI9 genetically and phenotypically with LB causing meningitis, increased inflammatory infiltrates and neuropathology *in vivo* and lytic replication *in vitro*, (25, 26, 33, 34). To define determinants of LB neurovirulence and mechanisms for attenuating LB, we generated an LB reverse genetics system. The robust CPER system developed permits the rapid generation of recLB and recLB EDIII mutants, and the evaluation of recombinant POWVs in a lethal murine model. Given the lethality of LI9 infection in aged mice, we inoculated 50-week-old mice with LB and evaluated lethal disease and histopathology. B6 mice succumbed to LB (79%) between 8 and 16 dpi with signs of lethargy, weakness, severe weight loss, and failure to self-right 8-11 dpi and severe neurological symptoms (eg. hindlimb paralysis) after 12 dpi. Overall, LB infected brains show evidence of meningoencephalitis, characterized by profound gliosis, perivascular cuffing, and cellular infiltration, that except for gliosis, are absent from previous reports of LI9 infection. Concurrent with high levels of viral RNA and gliosis in the LB infected CNS was the induction of interferon stimulated genes, proinflammatory cytokines and chemokines in LB infected mice. Further immunohistochemical analysis revealed POWV LB envelope protein immunostaining primarily to the cerebral cortex, hippocampus, caudatepatemun, and thalamus regions of the CNS. Unexpectedly, Iba1^+^ immunostaining of activated microglia/macrophages was primarily found at discrete CNS sites from antigen staining, and in contrast concentrated in foci within the midbrain, cerebellum, pons and medulla (Fig 5 and 6). This differs substantially from the staining of LI9 infected CNS where nearly uniform Iba^+^ staining was observed throughout the CNS (26). Whether these findings reflect LB inhibition of microglial/macrophage activation, or microglial cell clearance of LB in discrete CNS locations, remains to be investigated.

At 30 dpi, the CNS of LB infected survivors reveals high levels of Iba1^+^ staining which suggests persistent microglia/macrophage activation in mice surviving LB infection. In contrast, viral RNA levels and envelope antigen within the CNS of survivors are undetectable, suggesting the apparent clearance of virus and viral RNA from the CNS (Fig 4A and S2). Similarly, the CNS of LI9 survivors (30 dpi) was found to have sustained Iba1^+^ staining along with CD4 and CD8 T cell infiltrates (16). These findings suggest that following POWV clearance from the CNS, persistent CNS inflammation may contribute to long-term neurologic sequelae observed in POWV survivors (11, 50). However, a more detailed analysis of CNS responses, inflammation and immune cell recruitment is required to delineate potential causes of long-term CNS degeneration in survivors.

Strain LB was isolated from the brain of a fatal encephalitic patient, serially passaged i.c. in mouse brains and subsequently passaged in tissue culture (4, 28, 29). The overall impact of serial passage in mouse brains on systemic spread and neurovirulence has not been investigated. Our histological analysis of LB in the infected mouse CNS reflects findings from the original patient autopsy, where gliosis and perivascular infiltration was observed throughout the brain and specifically noted in the cortex, pons, and medulla (4). Other reports have shown LB infection of 4- and 6-week-old mice results in more widespread infection, including the cerebellum, thalamus, cortex, and brainstem (28, 29, 33). Consistent with our findings, one study reported microgliosis within the brainstem and cerebellum of LB infected mice, as a potential cause of immune mediated neuropathology in mice (29).

Changes in envelope proteins and EDIII have been associated with altered Flavivirus virulence. Structural analysis of tick borne encephalitis virus (TBEV) revealed a salt bridge between residues D308 and K311 in EDIII (37), and it was subsequently shown that mutations in these residues were partially attenuating (31). In Langat virus, D308A mutations were shown to cause a significant delay in lethality in infected SCIDs mice (51), while mutating D308N in Louping ill virus led to decreased neurovirulence (52). A novel case of fatal YFV encephalitis further suggested a role for EDIII K303Q mutations (311 equivalent in TBEV) in converting an attenuated YFV to neurovirulence (32).

Prior analysis of LI9 infected mice established that EDIII D308N mutations completely abolish LI9 lethality and infection of the CNS (27). LB differs from LI9 by one residue within an otherwise highly conserved region within EDIII, suggesting that both residues D308N and A310T could similarly impact LB neurovirulence and lethality. LB reverse genetics was used to make single D308N and double D308N/A310T LB mutants that were tested for their ability to cause lethal disease. Our results reveal that the LB-D308N mutant was partially attenuated and caused delayed lethality in only 33% of mice, with moribund mice having varying degrees of hindlimb paralysis. In contrast, the LB-D308N/A310T mutant that reflects residues in attenuated LI9P was fully attenuated in aged mice with no signs of neurological symptoms, weight loss, CNS pathology, gliosis or viral RNA in the CNS. In contrast to LB, LB-D308N/A310T infection failed to induce interferon stimulated genes, cytokines or chemokines in the CNS of infected mice. Immunohistochemical analysis of the infected CNS further substantiated these findings and revealed a total lack of POWV antigen in the CNS of LB-D308N/A310T infected mice at 10 or 30 dpi. Our findings reveal that POWV residues D308N and A310T play a critical role in POWV neurovirulence and lethality. Overall, LB-D308N/A310T infection resulted in the absence of microglial activation, CNS pathology, immune cell infiltrates and viral RNA and antigen in the CNS. These findings suggests that EDIII mutations D308N and A310T render LB unable to enter and infect the CNS, leading to complete attenuation of neurovirulence. However, more analysis is needed to understand the interactions of EDIII residues in preventing POWV neuroinvasion and whether differences reflect novel viral clearance, receptor attachment or cell tropism of EDIII mutants as mechanisms of attenuation and targets for rational vaccine development.

Our analysis of LB infected murine brains reveals discrete patterns of gliosis independent of POWV antigen staining that suggests potential viral clearance and immunoregulation in the LB infected CNS contribute to pathogenesis. Although these studies focused on envelope protein EDIII residues in LB neurovirulence, additional comparisons of LB and LI9 infections are needed to define determinants of lytic infection, peripheral spread, POWV receptors and the function of EDIII structures in the 5-fold axis of symmetry in POWV neuroinvasion. Collectively, our findings demonstrate that EDIII residues determine the neurovirulence and lethality of both lineage I and lineage II POWVs and reveal residues D308N and A310T as critical for POWV directed neuroinvasion and lethality.

## Materials and Methods

### Cells and Viruses

VeroE6 (ATCC CRL 1586) and BHK-21 [C-13] (ATCC CCL-10) were grown in Dulbecco’s modified Eagle’s medium at 37°C in 5% CO_2_. WT POWV-LB was obtained from the ATCC (NC003687). POWVs were absorbed to 70% confluent VeroE6 cell monolayers. After 2 hours, monolayers were PBS washed and grown in DMEM-2% FBS. POWV titers were determined by focus forming assay. Briefly, POWVs were serially diluted on VeroE6 and 30 hpi infected foci were quantified by immunoperoxidase staining with anti-POWV hyperimmune mouse ascites fluid (HMAF; 1:4,000; ATCC). All work with infectious POWVs was conducted in a certified BSL3 containment facility. Sequencing was performed by the SBU Genomics facility as previously described (35).

### Biosafety and security

Murine studies were conducted following institutional guidelines with approved experimental protocols and under the supervision of the Institutional Biosafety and Institutional Animal Care and Use Committees at SBU. Animals were managed by the SBU Division of Laboratory Animal Resources, accredited by the American Association for Accreditation of Laboratory Animal Care and DHHS, and maintained in accordance with the Animal Welfare Act and DHHS “Guide for the Care and Use of Laboratory Animals.” Veterinary care was directed by full-time resident veterinarians accredited by the American College of Laboratory Animal Medicine. POWV murine infection experiments were performed in an animal biosafety level 3 facility (The Laboratory of Comparative Medicine, Stony Brook University).

### Murine inoculation

C57BL/6J (B6) mice (50-week-old; Jackson Laboratory) were anesthetized via intraperitoneal injection of 100 mg of ketamine and 10 mg of xylazine per kilogram of body weight. Mice footpads were subcutaneously infected with 2×10^3^ FFU of POWV or mock infected with PBS in 20μl. Mice were weighed daily and checked for signs of neurological disease. Mice that reached humane endpoint criteria were euthanized from 1 to 30 days post infection. Criteria included non-responsiveness and severe neurological symptoms (hindlimb paralysis, ataxia, failure to self-right). Murine lethality studies were replicated in two independent experiments.

### RNA extraction and qRT-PCR

Brains were harvested from POWV infected moribund mice and homogenized in TRIzol LS Reagent (Invitrogen). RNA was isolated according to manufacturer’s protocol and purified by Monarch RNA Cleanup Kit (NEB T2030L). RNA quantification was performed on a Nanodrop Spectrophotometer 2000. cDNA synthesis was performed using the random hexamer priming of a ProtoScript First Strand cDNA Synthesis Kit (New England Biolabs) as previously described. qRT-PCR analysis of cDNA was used to determine viral loads with LB NS5-specific primers (Table 1). Each transcript was analyzed with technical duplicates using PerfeCTa SYBR green SuperMix (Quanta Biosciences) on a CFX opus 96 (Biorad). RNA levels were compared to a standard curve of serially diluted POWV-LB cDNA to determine FFU equivalents equating CT values to FFUs. To quantitate ISG and inflammatory transcript levels in POWV-infected brains, primers were designed with a 60°C annealing temperature using NCBI gene database (Table 1). Transcripts were analyzed in technical duplicate from n=3 mock or n=3 POWV infected mice. Responses were normalized to internal GAPDH levels, and the threshold cycle (2-ΔΔCT) method was used to determine the fold induction relative to controls.

### Histopathology

Harvested brains were fixed in 10% neutral buffered formalin for 7 days, dehydrated in 70% ethanol for 24 hours, and then paraffin embedded. The formalin-fixed paraffin embedded brain tissues were sagittally sectioned (4 µm thickness) and hematoxylin and eosin stained by Histowiz Inc. Immunohistochemical staining was performed by Histowiz with anti-Iba1 antibody (Wako 019–19741) to identify microglia and anti-POWV Env (GeneTex, GTX640340) to detect LB envelope protein. ImageJ was used to quantify Iba1^+^ pixel intensity percent area for 10 regions of POWV infected or mock midbrains. Qupath software (https://qupath.github.io) was used for tissue section analysis.

### Neutralizing antibody assay

C57BL/6 50-week-old mice were s.c. footpad inoculated with 2 × 10^3^ FFU of POWV or mock infected with PBS. At 28 days post infection, blood was collected from each mouse retro-orbitally under anesthesia (intraperitoneal injection of 100 mg of ketamine and 10 mg of xylazine per kilogram of body weight) and sera was isolated (Z-gel microtubes, Avantor). Neutralizing antibodies in the sera were assessed by focus reduction neutralization assays as previously described(^26, 27^). Complement proteins in the sera were inactivated at 56°C for 30 min. Then, sera were two-fold serially diluted and added to 500 FFU of POWV-LB for 1 hour. POWV was then absorbed to VeroE6 for 2 hours, and 36 hpi infected cell foci were detected by immunoperoxidase staining with anti-POWV HMAF. Serum dilutions required to reduce POWV foci by 50% were determined and presented as the neutralizing antibody titer.

### PCR amplification and CPER reverse genetics

Five POWV-LB (NCBI: NC003687) or LB mutant fragments were amplified from viral cDNA using high-fidelity Phusion polymerase (NEB) and corresponding paired primers that have a complementary 26 nucleotide overlap. An additional UTR linker fragment was generated from plasmid pMiniT containing the CMVd2 promoter, the first and the last 26 nts of the POWV-LI9 sequence, an HDVr, and SV40 polyadenylation site. POWV cDNA was amplified into six individual POWV DNA fragments: initial denaturation 98°C for 30 s; 32 cycles of 98°C for 20 s, 60°C for 30 s, and 72°C with 30 s per kb; and a final extension at 72°C for 5 min. Fragments were gel purified and isolated using Monarch DNA Gel Extraction kit (New England Biolabs). DNA fragments 1-5 and linker (0.09 pmol each) were CPER amplified in a 25-µL reaction containing 200 µM of dNTPs, Phusion polymerase GC reaction buffer, and 0.5 µL Phusion polymerase (New England Biolabs). The following cycling conditions were used: initial denaturation 98°C for 2 min; 18 cycles of 98°C for 30 s, 60°C for 30 s, and 72°C for 7 min; and a final extension at 72°C for 12 min.

To generate recombinant LB-D308N and LB-D308N/A310T, pMiniT plasmids were subjected to site-directed mutagenesis with Phusion polymerase and mutagenic primers (Table 1) to generate EDIII protein mutations at residue 308 and 310. Mutated plasmids were transformed into competent E coli Stable cells (NEB). Colonies were grown overnight and plasmids in single colonies were screened, sequenced and grown in LB broth and Miniprep (Qiagen) purified for fragment amplification.

### CEPR transfection and viral rescue

CEPR reactions were gel purified (Monarch Gel Extraction Kit, NEB) and eluted in nuclease free H2O and Lipofectamine 3000 (Invitrogen) transfected into BHK-21 cells in 6-well plates. Supernatants were harvested 3 days post transfection and amplified in VeroE6 cells to generate viral stocks. Total RNA from VeroE6 cell infections with LB, LB-D308N and LB-D308N/A310T was extracted and purified using RNeasy (Qiagen). cDNA synthesis was performed as above using random hexamer primers and cDNA was Sanger sequenced by the SBU Genomics Core.

### Western Blot Analysis

VeroE6 cells were infected with LB or LB-D308N/A310T (MOI, 0.1), and cells were harvested at 7 dpi in lysis buffer containing 1% NP-40 (150 mM NaCl, 50 mM Tris-Cl, 10% glycerol, 2 mM EDTA, 10 nM sodium fluoride, 2.5 mM sodium pyrophosphate, 2 mM sodium orthovanadate, 10 mM β glycerophosphate) with 1× protease inhibitor cocktail (Sigma). Total protein levels were determined in a bicinchoninic acid assay (Thermo Scientific) and 40 µg of protein was resolved by SDS-10% polyacrylamide gel electrophoresis. Proteins were then transferred to nitrocellulose, blocked in 5% milk, and incubated with rabbit anti-POWV EDIII antibody (1:4000). Protein was detected using HRP-conjugated anti-rabbit secondary antibody (1:4000) (Amersham) and Luminata Forte Western HRP substrate (Millipore).

### Statistical Analysis

Statistics were performed using Prism 10 software (GraphPad Software, Inc.). Statistical analysis of each experiment is described in its corresponding figure legend. The lethality evaluation of recombinant POWVs was replicated in two independent experiments. Kaplan-Meier curves were analyzed by a log rank test. Histopathology and IHC quantification were derived by Qpath and ImageJ analysis. Each brain was analyzed in 10 different locations compared across groups and *P* values of less than 0.05 were considered statistically significant.

## Data Availability Statement

The methods underlying this study are available in an online supplement at the authors’ institutional website and by request.

## Acknowledgements

We thank Megan Valenti, Ketaki Ganti, Catherine Finnerty and Varvara Kirillov for helpful discussions and feedback; Jorge Benach for continuing discussions of tick-borne diseases and neuropathogenesis and Aisling Byrne, Autumn Laird and Stella Tsirka for discussions of microglial CNS responses.

---

**Figure S1.**
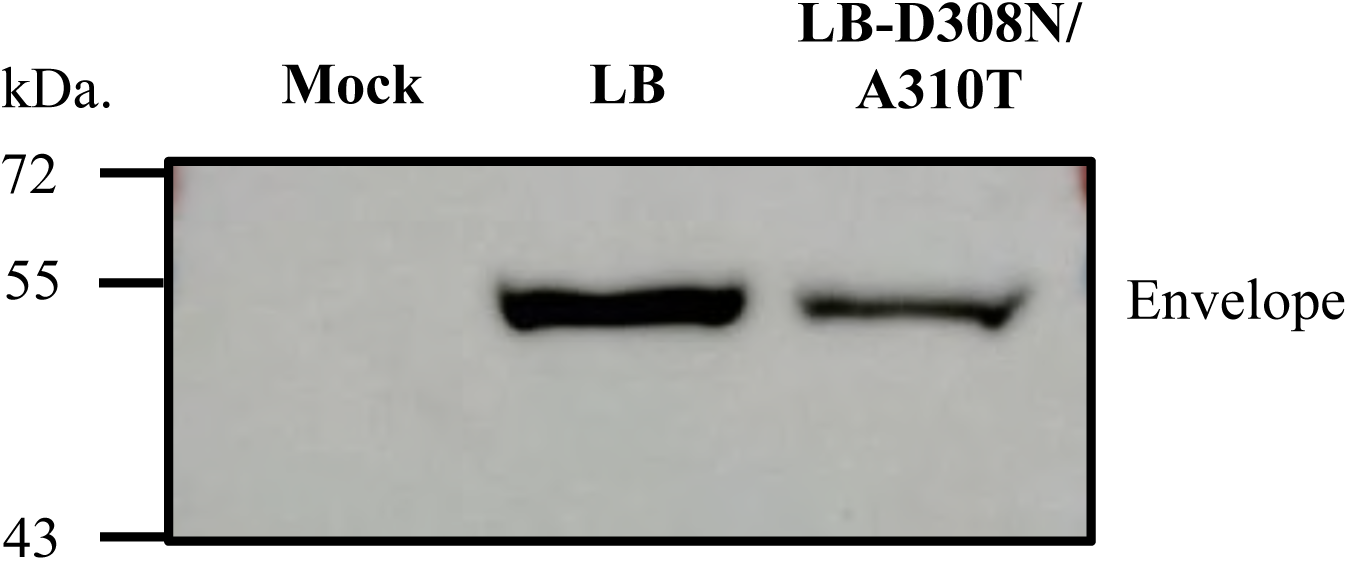
Analysis of Envelope Proteins from WT and NxT Added LB-D308N/A310T Mutant. VeroE6 Cells were infected with LB and LB-D308N/A310T (MOI 0.1) or mock infected. Cell lysates were harvested 7 dpi and 40 µg of total protein was analyzed by SDS-PAGE and Western Blotting with anti-POWV EDIII antibody (1:4000). (Env= POWV envelope protein).

**Figure S2.**
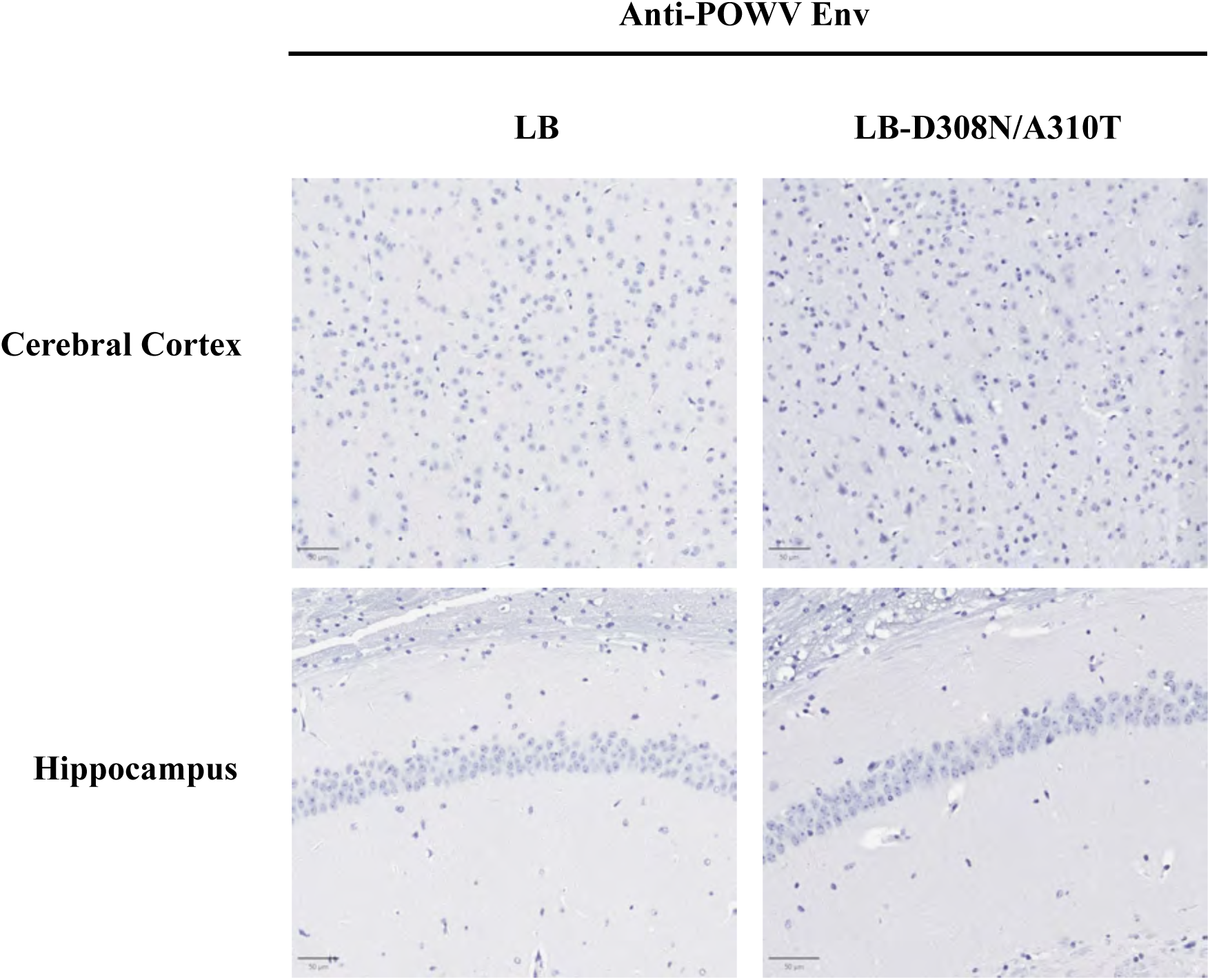
Analysis of the CNS of mice surviving LB and LB-D308N/A310T infection. (30 dpi). Surviving mice infected with LB (N=3) and LB-D308N/A310T (N=3) were euthanized at 30 dpi, brains were harvested, and immunostained with anti-POWV envelope protein antibodies. Representative staining of the hippocampus and cerebral cortex from WT and LB-D308N/A310 infected mice show the absence of POWV antigen in CNS.

## References

1. Pierson TC, Diamond MS. 2020. The continued threat of emerging flaviviruses. Nature Microbiology 5:796–812.

2. Kemenesi G, Bányai K. 2018. Tick-Borne Flaviviruses, with a Focus on Powassan Virus. Clinical Microbiology Reviews 32:10.1128/cmr.00106–17.

3. Sanchez-Vicente S, Tagliafierro T, Coleman JL, Benach JL, Tokarz R. 2019. Polymicrobial Nature of Tick-Borne Diseases. mBio 10.

4. Mc LD, Donohue WL. 1959. Powassan virus: isolation of virus from a fatal case of encephalitis. Can Med Assoc J 80:708–11.

5. El Khoury M, Camargo J, White J, Backenson B, Dupuis A, Escuyer K, Kramer L, St. George K, Chatterjee D, Prusinski M, Wormser G, Wong S. 2013. Potential Role of Deer Tick Virus in Powassan Encephalitis Cases in Lyme Disease–endemic Areas of New York, USA. Emerging Infectious Disease journal 19:1926.

6. Robich RM, Cosenza DS, Elias SP, Henderson EF, Lubelczyk CB, Welch M, Smith RP. 2019. Prevalence and Genetic Characterization of Deer Tick Virus (Powassan Virus, Lineage II) in Ixodes scapularis Ticks Collected in Maine. Am J Trop Med Hyg 101:467–471.

7. CDC. 2026. Powassan virus: historic data (2004–2024). Accessed 26 Jan.

8. Padda H, Huang CYH, Grimm K, Biggerstaff B, Ledermann J, Raetz J, Boroughs K, Mossel E, Martin S, Lehman J, Townsend R, Krysztof D, Saá P, Dinh ETN, Stobierski MG, Esponda-Morrison B, Wolujewicz KA, Osborne M, Brown C, Hopkins B, Schiffman E, Garvin A, Lee X, Osborn R, Wozniak R, Brault A, Basavaraju S, Stramer S, Staples JE, Gould C. 2025. Powassan and Eastern Equine Encephalitis Virus Seroprevalence in Endemic Areas, United States, 2019–2020. Emerging Infectious Disease journal 31:929.

9. Ebel GD, Kramer LD. 2004. Short report: duration of tick attachment required for transmission of powassan virus by deer ticks. Am J Trop Med Hyg 71:268–71.

10. Feder HM, Telford S, Goethert HK, Wormser GP. 2021. Powassan Virus Encephalitis Following Brief Attachment of Connecticut Deer Ticks. Clin Infect Dis 73:e2350–e2354.

11. Normandin E, Solomon IH, Zamirpour S, Lemieux J, Freije CA, Mukerji SS, Tomkins-Tinch C, Park D, Sabeti PC, Piantadosi A. 2020. Powassan Virus Neuropathology and Genomic Diversity in Patients With Fatal Encephalitis. Open Forum Infect Dis 7:ofaa392.

12. Burgos AF, Harford S, Kohli R. 2025. Emerging Threat: A Case of Neuroinvasive Powassan Virus Infection. Annals of Internal Medicine: Clinical Cases 4:e250097.

13. Telford SR, Piantadosi AL. 2023. Powassan virus persistence after acute infection. mBio 14:e0071223.

14. Chambers TJ, Hahn CS, Galler R, Rice CM. 1990. Flavivirus genome organization, expression, and replication. Annu Rev Microbiol 44:649–88.

15. Füzik T, Formanová P, Růžek D, Yoshii K, Niedrig M, Plevka P. 2018. Structure of tick-borne encephalitis virus and its neutralization by a monoclonal antibody. Nature Communications 9:436.

16. Mukhopadhyay S, Kim B-S, Chipman PR, Rossmann MG, Kuhn RJ. 2003. Structure of West Nile Virus. Science 302:248–248.

17. Das S, Narayanan A, Wang A, Moustafa IM, Cho SH, Mitzel D, Jose J, Hafenstein SL. 2025. Atomic-resolution structure of a chimeric Powassan tick-borne flavivirus. Science Advances 11:eadw7700.

18. Li C, Zhang L-y, Sun M-x, Li P-p, Huang L, Wei J-c, Yao Y-l, Isahg H, Chen P-y, Mao X. 2012. Inhibition of Japanese encephalitis virus entry into the cells by the envelope glycoprotein domain III (EDIII) and the loop3 peptide derived from EDIII. Antiviral Research 94:179–183.

19. Mertinková P, Mochnáčová E, Bhide K, Kulkarni A, Tkáčová Z, Hruškovicová J, Bhide M. 2021. Development of peptides targeting receptor binding site of the envelope glycoprotein to contain the West Nile virus infection. Scientific Reports 11:20131.

20. Kellman EM, Offerdahl DK, Melik W, Bloom ME. 2018. Viral Determinants of Virulence in Tick-Borne Flaviviruses. Viruses 10:329.

21. Beasley DWC, Suderman MT, Holbrook MR, Barrett ADT. 2001. Nucleotide sequencing and serological evidence that the recently recognized deer tick virus is a genotype of Powassan virus. Virus Research 79:81–89.

22. Kuno G, Artsob H, Karabatsos N, Tsuchiya KR, Chang GJ. 2001. Genomic sequencing of deer tick virus and phylogeny of powassan-related viruses of North America. Am J Trop Med Hyg 65:671–6.

23. Ebel GD. 2010. Update on Powassan Virus: Emergence of a North American Tick-Borne Flavivirus. Annual Review of Entomology 55:95–110.

24. Vogels Chantal BF, Brackney Doug E, Dupuis Alan P, Robich Rebecca M, Fauver Joseph R, Brito Anderson F, Williams Scott C, Anderson John F, Lubelczyk Charles B, Lange Rachel E, Prusinski Melissa A, Kramer Laura D, Gangloff-Kaufmann Jody L, Goodman Laura B, Baele G, Smith Robert P, Armstrong Philip M, Ciota Alexander T, Dellicour S, Grubaugh Nathan D. 2023. Phylogeographic reconstruction of the emergence and spread of Powassan virus in the northeastern United States. Proceedings of the National Academy of Sciences 120:e2218012120.

25. Conde JN, Sanchez-Vicente S, Saladino N, Gorbunova EE, Schutt WR, Mladinich MC, Himmler GE, Benach J, Kim HK, Mackow ER. 2022. Powassan Viruses Spread Cell to Cell during Direct Isolation from Ixodes Ticks and Persistently Infect Human Brain Endothelial Cells and Pericytes. J Virol 96:e0168221.

26. Mladinich MC, Himmler GE, Conde JN, Gorbunova EE, Schutt WR, Sarkar S, Tsirka SA, Kim HK, Mackow ER. 2024. Age-dependent Powassan virus lethality is linked to glial cell activation and divergent neuroinflammatory cytokine responses in a murine model. J Virol 98:e0056024.

27. Himmler GE, Mladinich MC, Conde JN, Gorbunova EE, Lindner MR, Kim HK, Mackow ER. 2025. Passage-attenuated Powassan virus LI9P protects mice from lethal LI9 challenge and links envelope residue D308 to neurovirulence. mBio 16:e0006525.

28. Santos RI, Hermance ME, Gelman BB, Thangamani S. 2016. Spinal Cord Ventral Horns and Lymphoid Organ Involvement in Powassan Virus Infection in a Mouse Model. Viruses 8:220.

29. Reynolds ES, Hart CE, Nelson JT, Marzullo BJ, Esterly AT, Paine DN, Crooker J, Massa PT, Thangamani S. 2024. Comparative Pathogenesis of Two Lineages of Powassan Virus Reveals Distinct Clinical Outcome, Neuropathology, and Inflammation. Viruses 16:820.

30. Ryman KD, Ledger TN, Campbell GA, Watowich SJ, Barrett AD. 1998. Mutation in a 17D-204 vaccine substrain-specific envelope protein epitope alters the pathogenesis of yellow fever virus in mice. Virology 244:59–65.

31. Mandl CW, Allison SL, Holzmann H, Meixner T, Heinz FX. 2000. Attenuation of tick-borne encephalitis virus by structure-based site-specific mutagenesis of a putative flavivirus receptor binding site. J Virol 74:9601–9.

32. Jennings AD, Gibson CA, Miller BR, Mathews JH, Mitchell CJ, Roehrig JT, Wood DJ, Taffs F, Sil BK, Whitby SN, et al. 1994. Analysis of a yellow fever virus isolated from a fatal case of vaccine-associated human encephalitis. J Infect Dis 169:512–8.

33. Holbrook MR, Aronson JF, Campbell GA, Jones S, Feldmann H, Barrett ADT. 2005. An Animal Model for the Tickborne Flavivirus—Omsk Hemorrhagic Fever Virus. The Journal of Infectious Diseases 191:100–108.

34. Mlera L, Meade-White K, Saturday G, Scott D, Bloom ME. 2017. Modeling Powassan virus infection in Peromyscus leucopus, a natural host. PLOS Neglected Tropical Diseases 11:e0005346.

35. Conde JN, Himmler GE, Mladinich MC, Setoh YX, Amarilla AA, Schutt WR, Saladino N, Gorbunova EE, Salamango DJ, Benach J, Kim HK, Mackow ER. 2023. Establishment of a CPER reverse genetics system for Powassan virus defines attenuating NS1 glycosylation sites and an infectious NS1-GFP11 reporter virus. mBio 14:e0138823.

36. Malonis RJ, Georgiev GI, Haslwanter D, VanBlargan LA, Fallon G, Vergnolle O, Cahill SM, Harris R, Cowburn D, Chandran K, Diamond MS, Lai JR. 2022. A Powassan virus domain III nanoparticle immunogen elicits neutralizing and protective antibodies in mice. PLoS Pathog 18:e1010573.

37. Rey FA, Heinz FX, Mandl C, Kunz C, Harrison SC. 1995. The envelope glycoprotein from tick-borne encephalitis virus at 2 A resolution. Nature 375:291–8.

38. Rossier E, Harrison RJ, Lemieux B. 1974. A case of Powassan virus encephalitis. Can Med Assoc J 110:1173–4 passim.

39. Piantadosi A, Rubin DB, McQuillen DP, Hsu L, Lederer PA, Ashbaugh CD, Duffalo C, Duncan R, Thon J, Bhattacharyya S, Basgoz N, Feske SK, Lyons JL. 2015. Emerging Cases of Powassan Virus Encephalitis in New England: Clinical Presentation, Imaging, and Review of the Literature. Clinical Infectious Diseases 62:707–713.

40. Fagre AC LS, Staples JE, Lindsey N. 2023. West Nile Virus and Other Nationally Notifiable Arboviral Diseases — United States, 2021. MMWR Morb Mortal Wkly Rep 72.

41. Orozco S, Schmid MA, Parameswaran P, Lachica R, Henn MR, Beatty R, Harris E. 2012. Characterization of a model of lethal dengue virus 2 infection in C57BL/6 mice deficient in the alpha/beta interferon receptor. Journal of General Virology 93:2152–2157.

42. Meier KC, Gardner CL, Khoretonenko MV, Klimstra WB, Ryman KD. 2009. A Mouse Model for Studying Viscerotropic Disease Caused by Yellow Fever Virus Infection. PLOS Pathogens 5:e1000614.

43. Lee E, Lobigs M. 2008. E protein domain III determinants of yellow fever virus 17D vaccine strain enhance binding to glycosaminoglycans, impede virus spread, and attenuate virulence. J Virol 82:6024–33.

44. Zhang J, Chavez EC, Winkler M, Liu J, Carver S, Lin AE, Biswas A, Tamura T, Tseng A, Wang D, Benhamou A, O’ Connell AK, Matsuo M, Norton JE, Kenney D, Adamson B, Kleiner RE, Burwitz B, Crossland NA, Douam F, Ploss A. 2025. Amino acid changes in two viral proteins drive attenuation of the yellow fever 17D vaccine. Nature Microbiology 10:1902–1917.

45. Huang R, He Y, Zhang C, Luo Y, Chen C, Tan N, Ren Y, Xu K, Yuan L, Yang J. 2024. The mutation of Japanese encephalitis virus envelope protein residue 389 attenuates viral neuroinvasiveness. Virol J 21:128.

46. Hung JJ, Hsieh MT, Young MJ, Kao CL, King CC, Chang W. 2004. An external loop region of domain III of dengue virus type 2 envelope protein is involved in serotype-specific binding to mosquito but not mammalian cells. J Virol 78:378–88.

47. Mittler E, Tse AL, Tran PT, Florez C, Janer J, Varnaite R, Kasikci E, Mv VK, Loomis M, Christ W, Cazares E, Bakken RR, Martin CK, Zeng X, Raymond JL, Shahsavani M, Khanal S, Wilkinson ER, Oktavia RM, Slough MM, Haslwanter D, Han J, Berrigan J, Rosendal E, Kielian M, Manicassamy B, Överby AK, Falk A, Barba-Spaeth G, Rey FA, Klingström J, Gavathiotis E, Herbert AS, Chandran K, Gredmark-Russ S. 2025. LRP8 is a receptor for tick-borne encephalitis virus. Nature 646:945–952.

48. Chong Z, Hui S, Qiu X, Palakurty S, Sariol A, Kaszuba T, Nguyen MN, Li P, Raju S, Hall PD, Nelson CA, Baltazar-Perez I, Price DA, Rothlauf PW, Crowe JE, Whelan SPJ, Leung DW, Amarasinghe GK, Bailey AL, Fremont DH, Diamond MS. 2026. Multiple LDLR family members act as entry receptors for yellow fever virus. Nature 649:173–182.

49. Daskou M, Zaiss AK, Jeyachandran AV, Takano K-A, Kan RL, Paravastu R, Gerald E, Satheeshkumar N, Rios-Rodriguez J, Russell B, Brault AC, Bhaduri A, Garcia G, Jr., Morizono K, Arumugaswami V. 2025. Phosphatidylserine receptors TIM-1 and AXL mediate tick-borne Powassan virus entry. iScience 28.

50. Krow-Lucal ER, Lindsey NP, Fischer M, Hills SL. 2018. Powassan Virus Disease in the United States, 2006-2016. Vector Borne Zoonotic Dis 18:286–290.

51. Campbell MS, Pletnev AG. 2000. Infectious cDNA clones of Langat tick-borne flavivirus that differ from their parent in peripheral neurovirulence. Virology 269:225–37.

52. Jiang WR, Lowe A, Higgs S, Reid H, Gould EA. 1993. Single amino acid codon changes detected in louping ill virus antibody-resistant mutants with reduced neurovirulence. J Gen Virol 74 (Pt 5):931–5.

